# Architecture of a minimal signalling pathway explains the T cell response to a 1,000,000-fold variation in antigen affinity and dose

**DOI:** 10.1101/071878

**Authors:** Melissa Lever, Hong-Sheng Lim, Philipp Kruger, John Nguyen, Nicola Trendel, Enas Abu-Shah, Philip K. Maini, P. Anton van der Merwe, Omer Dushek

**Author notes:** These authors contributed equally (Melissa Lever, Hong-Sheng Lim, Philipp Kruger).

## Abstract

T cells must respond differently to antigens of varying affinity presented at different doses. Previous attempts to map pMHC affinity onto T cell responses have produced inconsistent patterns of responses preventing formulations of canonical models of T cell signalling. Here, a systematic analysis of T cell responses to 1,000,000-fold variations in both pMHC affinity and dose produced bell-shaped dose-response curves and different optimal pMHC affinities at different pMHC doses. Using sequential model rejection/identification algorithms, we identified a unique, minimal model of cellular signalling incorporating kinetic proofreading with limited signalling coupled to an incoherent feed forward loop (KPL-IFF), that reproduces these observations. We show that the KPL-IFF model correctly predicts the T cell response to antigen co-presentation. Our work offers a general approach for studying cellular signalling that does not require full details of biochemical pathways.

**Significance statement:** T cells initiate and regulate adaptive immune responses when their T cell antigen receptors recognise antigens. The T cell response is known to depend on the antigen affinity/dose but the precise relationship, and the mechanisms underlying it, are debated. To resolve the debate, we stimulated T cells with antigens spanning a 1,000,000-fold range in affinity/dose. We found that a different antigen (and hence different affinity) produced the largest T cell response at different doses. Using model identification algorithms, we report a simple mechanistic model that can predict the T cell response from the physiological low affinity regime into the high affinity regime applicable to therapeutic receptors.

## Introduction

T cell activation is critical for initiating and regulating adaptive immunity (1). It proceeds when T cell receptors (TCRs) on the T cell surface bind to antigenic peptides loaded on major histocompatibility complexes (pMHCs). Binding of pMHC ligands to the TCR initiates a large signal transduction cascade that can lead to T cell activation as measured by functional responses such as proliferation, differentiation, target cell killing, and the production and secretion of effector cytokines. These responses critically depend on the pMHC affinity and dose. T cells are known to discriminate between normal and infected or cancerous cells based on differences in pMHC affinity (2, 3). It is also appreciated that the pMHC dose determines, for example, the peripheral induction of regulatory T cells (4, 5). Although the proteins that form the TCR-regulated signalling network have been identified (6, 7), it remains unclear how the architecture they form integrates the pMHC affinity and dose into T cell activation (8, 9).

Studies performed over the last two decades have focused on empirically mapping the relationship between pMHC affinity and T cell activation (5, 10–24). A number of studies have reported an optimal pMHC affinity for T cell activation but other studies have failed to observe the optimum. Interestingly, a subset of studies have suggested that the optimal pMHC affinity may be less pronounced at high pMHC doses (13, 22). The mechanism underlying an optimal pMHC affinity (or half-life) is proposed to be a trade-off between serial binding and kinetic proofreading but we have recently shown that this trade-off would lead to an optimal pMHC affinity at all pMHC doses (9).

An accurate model of T cell signalling pathways that can predict the T cell response to a broad range of antigen ligand affinity and dose is important not only to understand physiological T cell responses but also in the rational design of T cell based therapies (25). Engineered therapeutic TCRs and chimeric antigen receptors (CARs) have been produced to bind, for example, cancer antigens with high affinity (dissociation constants, *K*_D_, in the picomolar to nanomolar range). A specific example is the NY-ESO-1 cancer antigen, for which both high affinity TCRs and CARs have been produced (26, 27). The optimising of these therapies has focused, in part, on trying to determine the optimal receptor affinity for clinical efficacy (20, 22, 28–30) (reviewed in (31)).

A key challenge in the study of cellular signalling in general, and particularly in T cells, is the organisation of large amounts of molecular information into accurate mathematical models that can predict cellular responses (32). The reductionist approach has been to incorporate the known biochemistry into mathematical models, but it is well recognised that this relies on many assumptions (e.g. which proteins and interactions to include, their binding and reaction rate constants, concentrations, etc.) leading to models whose accuracy is difficult to determine (32, 33). This may, in part, explain why canonical models of T cell signalling have been elusive (Figure 1a). An alternative holistic approach is to infer models from experimental data without any prior assumptions (34–39) (e.g. of the known biochemistry).

In this work, we utilised the high affinity engineered c58c61 TCR that binds the NY-ESO-1 cancer antigen (27) to measure primary human T cell activation in response to a 1,000,000-fold variation in pMHC affinity and dose. We found bell-shaped dose-response curves with inhibition at high pMHC doses. Moreover, different pMHCs (and hence different affinities) produced the largest T cell response at different pMHC doses. Without making prior assumptions about the known biochemistry, we identified a unique and modular pathway architecture for cellular signalling that reproduced these observations: kinetic proofreading with limited signalling coupled to an incoherent feedforward motif (KPL-IFF). We show that the identified KPL-IFF model predicts the outcome of pMHC co-presentation experiments. These mechanistic insights force a revision of the serial binding and kinetic proofreading model for T cell activation. The revised KPL-IFF model now predicts T cell activation from the physiological low affinity regime into the high affinity regime applicable to therapeutic TCRs and CARs.

## Results

### T cell activation in response to a 1,000,000-fold variation in antigen affinity/dose

As a first step to identify a T cell signalling model (Figure 1a) we established a TCR/pMHC system with a large range of affinities by using the therapeutic affinity-matured c58c61 TCR that recognises a peptide derived from the cancer antigen NY-ESO-1 in complex with HLA-A*02:01 (*K*_D_ ~ 50 pM) (27). This TCR contains 14 amino acid substitutions (primarily at the contact interface) but maintains the same binding mode as the parental 1G4 TCR (40). Using single, double, and triple peptide mutations we produced a panel of 11 pMHCs that span a 1,000,000-fold range in affinity (Figure 1b-c and SI Appendix, Table S1, Figure S1). The observed changes in affinity were largely a result of changes in the off-rate *k*_off_.

We next transduced the c58c61 TCR into primary human CD8+ T cells and the Jurkat T cell line (SI Appendix, Figure S2a) before stimulating them with a 1,000,000-fold range of pMHC concentration. In the case of the primary T cells we measured the supernatant concentration of IFN-*γ* or MIP-1*β* after 4 hours of stimulation (Figure 1d-e and SI Appendix, Figure S2b). In the case of the Jurkat T cells we measured the supernatant concentration of IL-8 after 16 hours (Figure 1f and SI Appendix, Figure S2c) or the transcriptional activity of AP1/NFAT (SI Appendix, Figure S2d).

**Figure 1:**
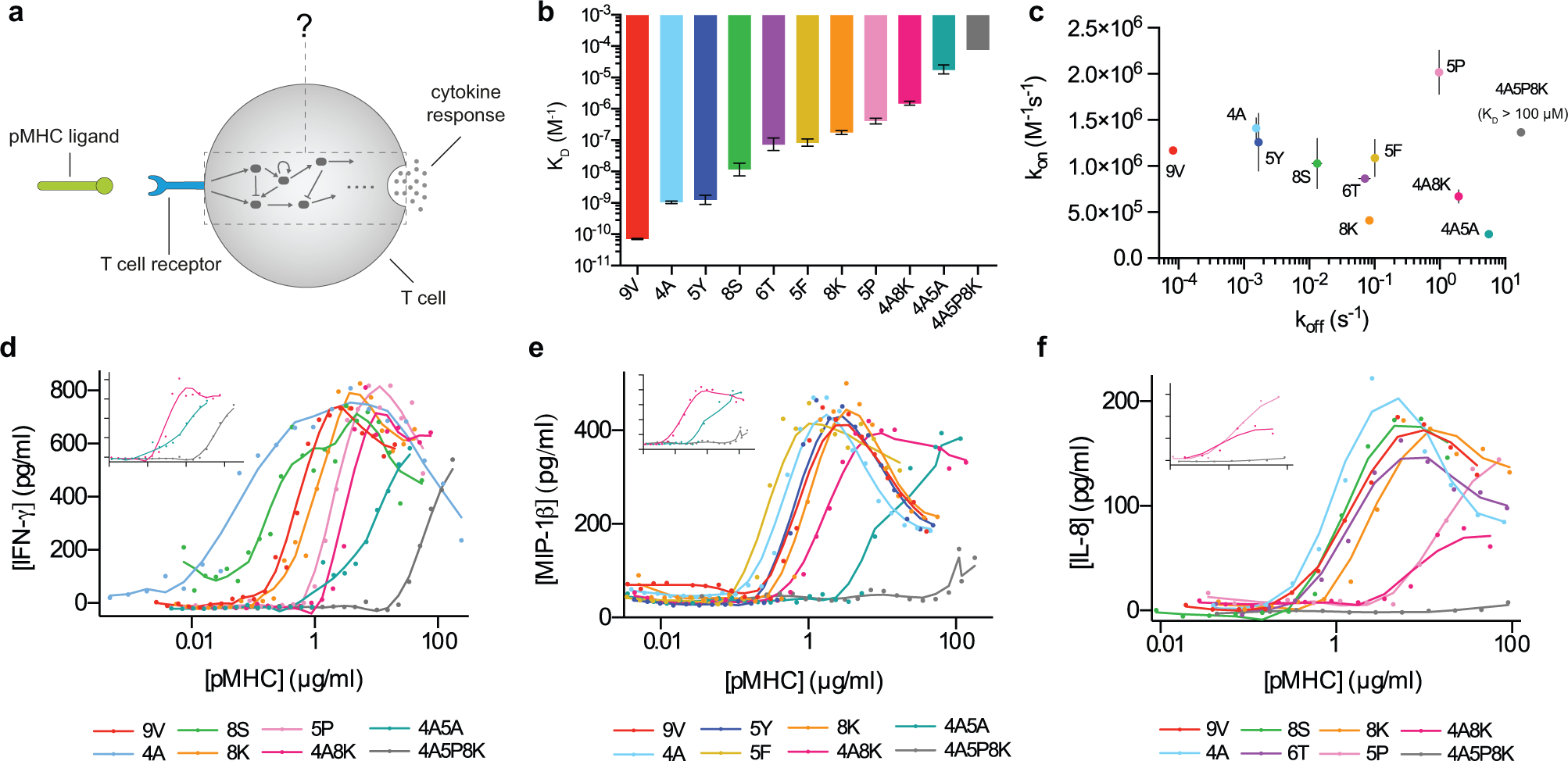
T cell activation in response to a 1,000,000-fold variation in pMHC affinity and dose. a) Schematic illustrating that the signalling architecture linking the TCR to cytokine production is unknown. b) Affinities (displayed as dissociation constants, *K*_D_) and c) kinetics of the c58c61 TCR interacting with 11 pMHC ligands determined using surface plasmon resonance (see also SI Appendix, Figure S1, Table S1). T cell activation as measured by supernatant d) IFN-*γ* and e) MIP-1*β* in primary T cells after 4 hours and f) IL-8 in Jurkat T cells after 16 hours transduced with the c58c61 TCR (inset shows the 3 lowest affinity ligands in each experiment). Ligand color scheme is identical across all panels. Additional data, including additional concentrations, additional ligands, pMHC immobilisation controls, and TCR expression levels, are summarised in SI Appendix, Figure S2.

Four striking features were observed. First, the dose-response appeared to exhibit a bell-shape with reduced cytokine production at high pMHC concentrations. This bell-shape was less pronounced or absent for low affinity ligands, which is consistent with published studies reporting a sigmoidal dose-response for low affinity ligands (14, 15, 17, 18, 21, 23, 41). Second, the peak amplitude of the bell-shaped dose-response was similar for pMHCs despite large differences in their affinities. The next two features describe the observation that the pMHC that produced the most cytokine was dose dependent. At higher concentrations (to the right of the peaks) there is an obvious intersection of the dose-response curves so that different pMHC ligands produce the most cytokine at different doses (e.g. 4A8K or 4A5A in Figure 1d-f). In contrast, at lower concentrations (to the left of the peaks) curve intersection is not apparent so that a single intermediate affinity pMHC produces the most cytokine. These four phenotypic features are summarised in Table 1.

**Table 1:**
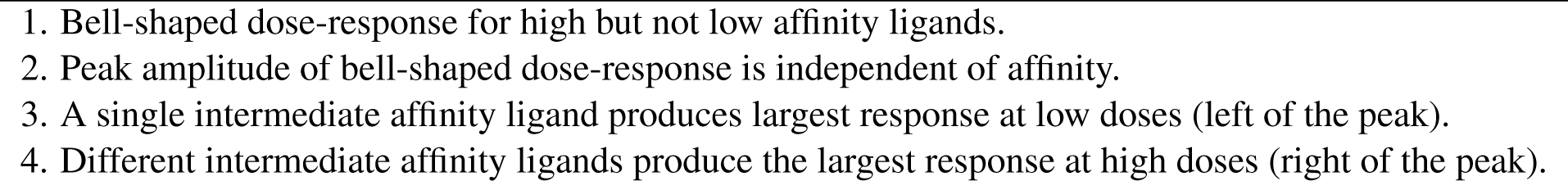
Phenotypic features of T cell activation

While a bell-shaped dose-response can be a result of activation induced cell death (42), this is unlikely to be the case here. We found that the dose-response for cytokine production appeared bell-shaped at all times with continued cytokine production to the right of the peak (Figure 2a). Moreover, direct detection of apoptosis by Annexin V binding revealed a maximum increase of only 10%, which is insufficient to explain the observed reduction in cytokine production (Figure 2b). Further, lower levels of cytokine observed to the right of the peak were associated with lower levels of Annexin V, which is inconsistent with the hypothesis that lower cytokine is a result of increased apoptosis at high pMHC doses. These observations suggested a reduced rate of cytokine production per cell at high pMHC doses, which we confirmed using single cell cytokine production in Jurkat T cells (Figure 2c) and in primary T cells (SI Appendix, Figure S2e).

We observed inter-donor variability that could not be explained by differential TCR expression or pMHC activity. For example, the pMHC that produced the largest response to the left of the peak varied between 4A, 5Y, and 8S (compare Figure 1d and SI Appendix, Figure S2b). However, the feature that a single pMHC of intermediate affinity produced the largest response to the left of the peak was consistent. Therefore, although the quantitative features of the data exhibited variability, we observed a high level of consistency for the key qualitative phenotypic features (Table 1). Similarly, we found that Jurkat T cells required a higher amount of antigen to produce cytokine but that the overall response exhibited the same qualitative phenotypic features observed in the primary T cells. Although these Jurkats express CD8α, their reduced sensitivity may be related to the absence of CD8β, which has previously been shown to increase antigen sensitivity (12).

### Sequential model rejection identifies the KPL-IFF model as sufficient to explain T cell responses

We next identified a T cell signalling network consistent with all key features by sequentially rejecting models. To do this, we tested models of increasing complexity starting with the simplest possible cellular mechanism, namely the occupancy model, which reduces cellular signalling to a single reaction (Figure 3a). In this model, the pMHC ligand binds to the TCR to form a complex that directly activates a protein *P* that is assumed to be linearly proportional to cytokine production. By examining the predicted dose-response for this model it is clear that the model is insufficient to explain the phenotypic features (e.g. it does not produce a bell-shaped dose-response, feature 1) and therefore we reject this model as a plausible model of T cell signalling.

Bell-shaped dose-response curves can be produced by incoherent feedforward motifs (Figure 3b), which are common architectures in transcriptional networks (43). In this model, the TCR-pMHC complex directly inhibits *P* and indirectly activates *P* (by activating *Y* which itself is able to activate P). The model can produce inhibition at high pMHC concentrations if the activatory pathway (through *Y*) saturates, allowing inhibition to dominate at the highest pMHC concentrations. The appearance of the bell-shape in this model can explain the observation that a different pMHC affinity produces the largest response to the right of the peak (feature 4). However, this model is also rejected because it produces a bell-shaped dose-response for all pMHC ligands independent of their affinity (feature 1).

**Figure 2:**
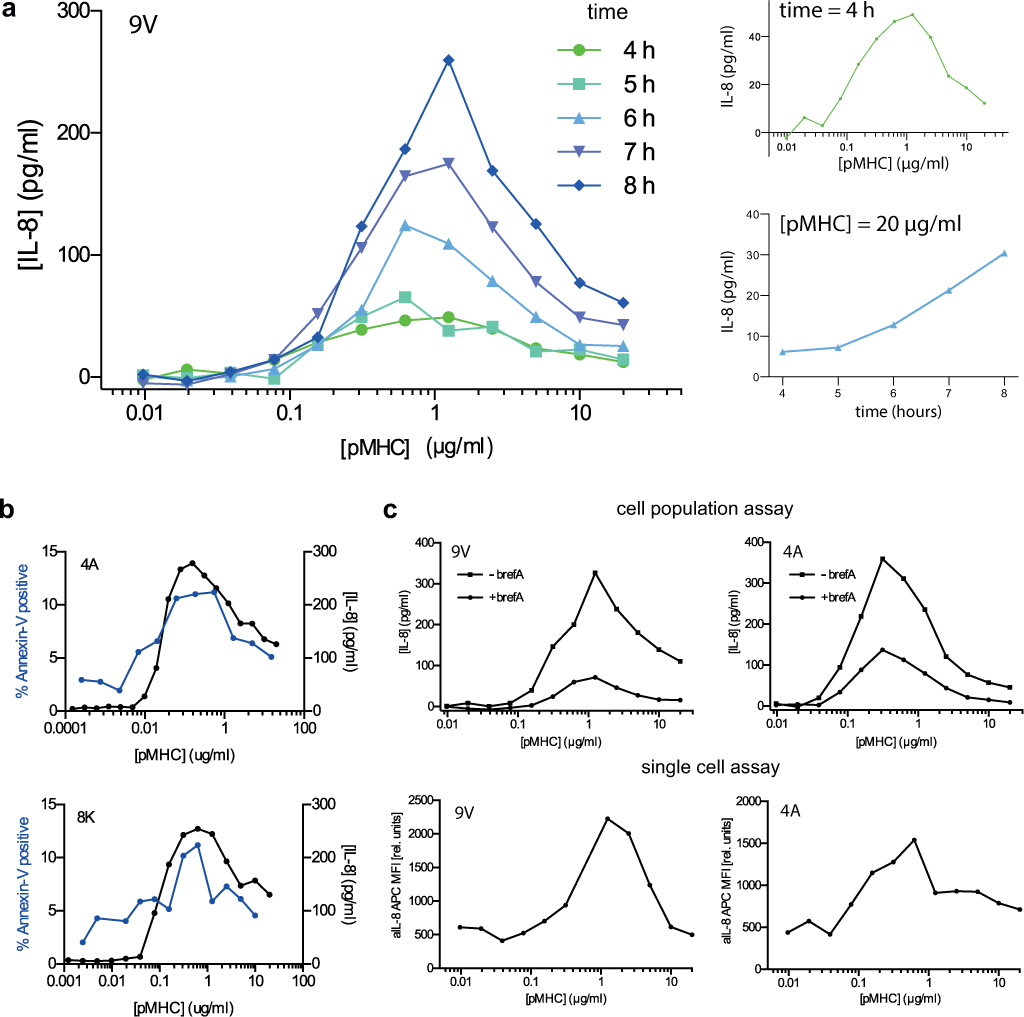
A bell-shaped dose-response is a consequence of reduced cytokine production at high pMHC concentrations. a) T cell activation dose-response curves at the indicated time points (left panel) highlighting the bell-shape at early times (4 hours) and continued cytokine production at the largest pMHC concentration (right two panels). b) Percent of T cells positive for Annexin V (blue, left axis) determined at the end of a 16 hour functional assay where the supernatant concentration of IL-8 was also determined (black, right axis). c) Comparison of supernatant IL-8 production at the population level (top) with the corresponding single cell IL-8 production by flow cytometry (bottom) at 16 hours. Brefeldin A was added to block cytokine secretion for the last 3 hours of the assay (reducing supernatant cytokine in the cell population assay). Jurkat T cells are used to generate all panels with the indicated pMHC ligands. See SI Appendix, Figure S2e for single cell cytokine production in primary CD8+ T cells.

Bell-shaped dose-response curves can be produced for high but not low affinity ligands by introducing kinetic proofreading (Figure 3c). In this model, the pMHC ligand does not trigger signalling immediately upon binding TCR but instead must remain bound until it becomes signalling competent (denoted as *C*_1_). This delay means that low affinity pMHCs (with faster *k*_off_) induce a lower maximal concentration of *C*_1_ than high affinity pMHC (see SI Appendix, Figure S3 for a plot of C_1_ for pMHC with different values of *k*_off_). If this is below the level at which *Y* saturates then inhibition at high pMHC concentrations will not be observed with low affinity pMHC. As expected, kinetic proofreading has improved antigen discrimination by dramatically decreasing the T cell response to low affinity pMHC. This model, however, is also rejected because it predicts that the highest affinity ligand will produce the largest response left of the peak in contrast to experimental observations (feature 3).

Introducing limited signalling into kinetic proofreading can produce an optimal affinity over a range of pMHC concentrations (9) (Figure 3d). In this model, activated TCR-pMHC complexes (*C*_1_) signal for a limited period of time before converting to a non-signalling state (*C*_2_) thereby introducing a penalty for pMHC that remain bound for long periods of time. This model is now able to explain all key features (Table 1) and we therefore accept the kinetic proofreading with limited signalling coupled to an incoherent feedforward loop (KPL-IFF) model as a plausible signalling model for T cell activation.

**Figure 3:**
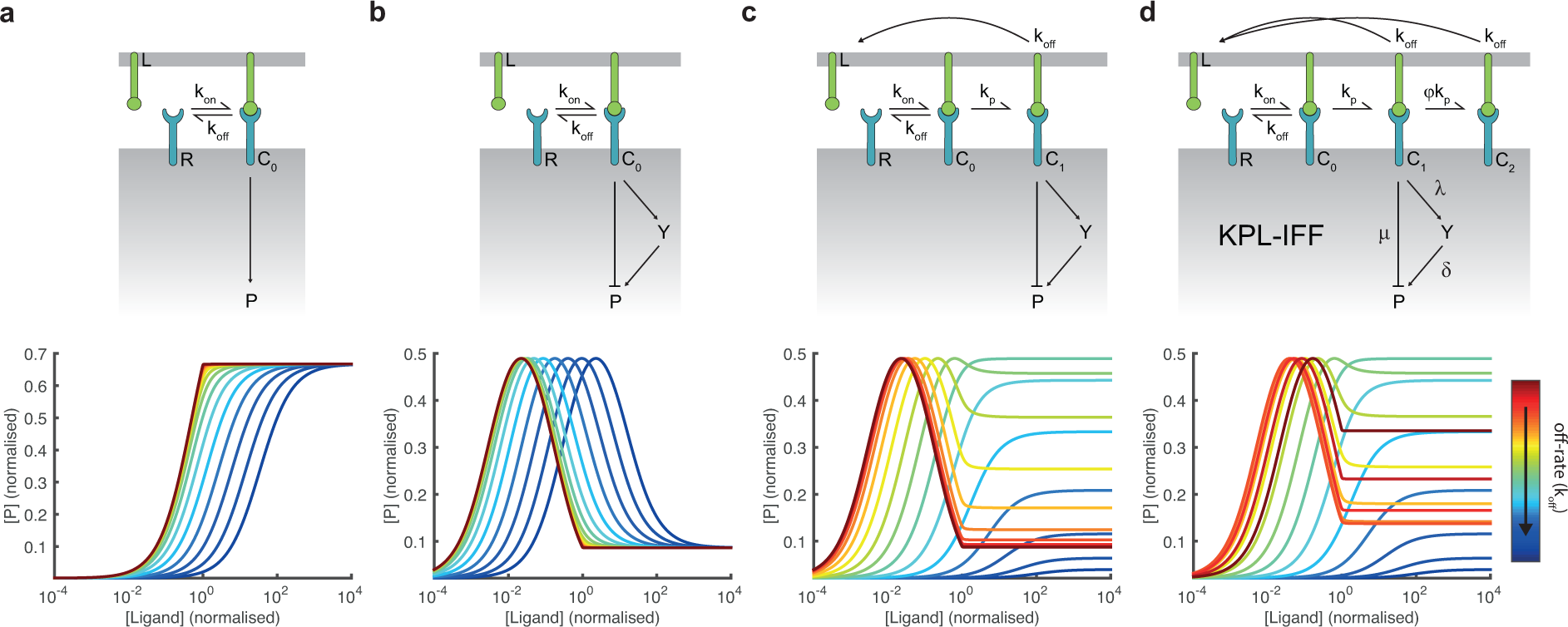
Sequential model rejection reveals that kinetic proofreading with limited signalling coupled to an incoherent feedforward loop can produce all phenotypic features (Table 1). The models considered, in order of increasing complexity, are a) occupancy, b) occupancy coupled to incoherent feedforward, c) kinetic proofreading coupled to incoherent feedforward, and d) kinetic proofreading with limited signalling coupled to an incoherent feedforward loop (KPL-IFF). All models include the reversible (serial) binding of pMHC ligands (*L*) to the TCR (*R*) to form complexes that can regulate the activation of a protein *P* that is taken to be a measure of T cell activation. See SI Appendix for computational details and Applet S1 for a tool that can be used to explore how the 5 parameters in the KPL-IFF model (*k*_p_, *ϕ*, *µ*, λ, and *δ*) modulate the predicted dose-response for antigens of different affinities.

### Systematic model identification confirms that the KPL-IFF model is unique

Having identified the KPL-IFF model as sufficient to explain all phenotypic features, we next determined whether other models, with a potentially different underlying mechanism, can also explain all phenotypic features.

We first systematically examined models of equal or lower complexity to the KPL-IFF model. To do this, we studied all combinations of 3 reaction arrows between *Y* and *P* and the 3 receptor states (Figure 4a). Of the 560 possible reaction networks (16 choose 3) only the 304 networks that contain a connection between the ligand and *P* were analysed. For each of these putative signalling networks, we performed an exhaustive search that included a dense parameter scan followed by optimisation of the 5 free parameters (see SI Appendix, Figure S4). The output of the analysis is a list of networks ordered by their ability to reproduce the phenotypic features (Movie S1). As expected, the first network to appear is the KPL-IFF model but, unexpectedly, the 303 subsequent networks were all unable to explain all phenotypic features.

We highlight 3 models from the network search that are inconsistent with the phenotypic features (Figure 4b-d). A mirrored model in which activation is direct but inhibition is indirect cannot produce a bell-shaped dose-response (feature 1) because activation cannot saturate (Figure 4b). A re-directed model where inhibition comes from an earlier complex not subjected to kinetic proofreading (Figure 4c) cannot explain a different optimal pMHC affinity to the right of the peak (feature 4) nor the observation that the peak response is similar for different affinity ligands (feature 2). Lastly, models without an incoherent feedforward loop but with negative feedback, although able to produce oscillations of *P* in time, cannot produce a bell-shaped dose-response (Figure 4d and Figure S5, see also SI Appendix for a mathematical proof).

**Figure 4:**
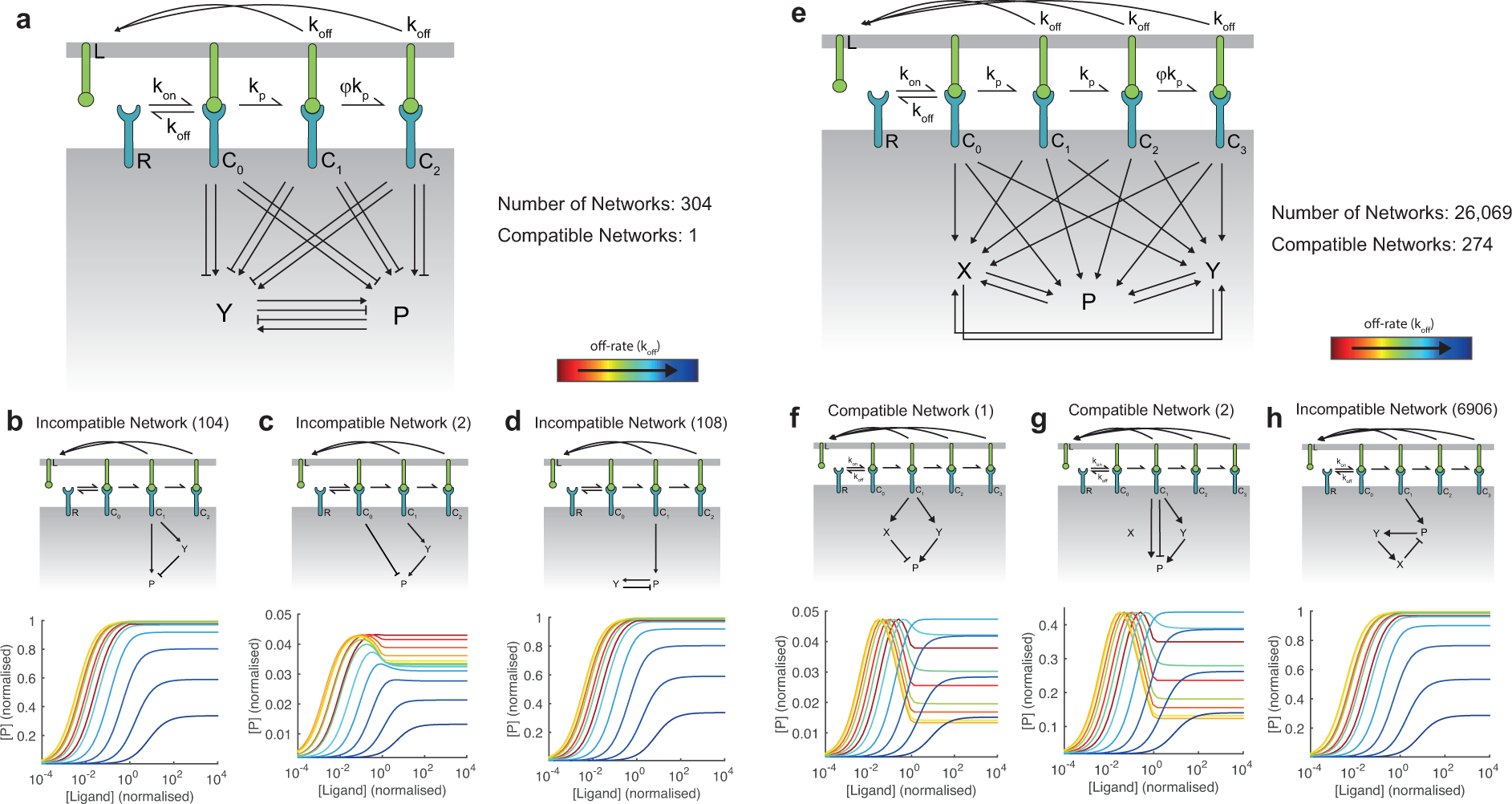
Systematic analyses of signalling models reveals that the KPL-IFF mechanism is unique. a) To determine if other models of equal (or lower) complexity to the KPL-IFF model (Figure 3D) are able to produce all phenotypic features we performed a systematic search of 304 network architectures with 3 reaction arrows between the receptor states (*C*_0_, *C*_1_, and *C*_2_), *Y*, and *P*. The only network architecture that is able to produce all phenotypic features is the KPL-IFF model (see Movie S1). Conversely, b) the mirrored KPL-IFF, c) the re-directed KPL-IFF, and D) negative feedback network architectures are unable to produce the phenotypic features. e) To determine if more complex models can reproduce the phenotypic features using mechanisms different from those invoked in the KPL-IFF model, we performed a systematic analysis of 26,069 network architectures with 4 reaction arrows between 4 receptor states and *Y*, *P*, and an additional node *X*. Both activation and inhibition are considered but for clarity only activation arrows are depicted. We found 274 networks compatible with all phenotypic features but all of these networks relied on the KPL-IFF mechanism (see Movie S2). f,g) Two representative compatible networks show that although the network is more complicated both rely on the KPL-IFF mechanism. h) As before, negative feedback in the absence of incoherent feedforward is unable to produce the phenotypic features. See SI Appendix for computational details.

To determine whether more complex models can explain all key features using different mechanisms, we performed the same systematic network analysis on models with 4 reaction arrows between *Y*, *P* and an additional node *X* and 4 receptor states (Figure 4e). A systematic analysis of the 26,069 networks with a connection between the ligand and *P* revealed 274 compatible networks (Movie S2). However, examining these 274 networks showed that the basic mechanism underlying all compatible networks was KPL-IFF. For example, the incoherent feedforward loop could involve indirect inhibition so long as inhibition saturates after activation as a function of ligand dose (Figure 4f) or it could involve both direct activation and inhibition with the net effect being inhibition (Figure 4g). As above, negative feedback could not produce bell-shaped dose-response curves (Figure 4h).

In summary, the KPL-IFF signalling network (Figure 3d) is sufficient to explain all phenotypic features. Given that the systematic analyses implicitly include simpler models (e.g. by allowing for the magnitude of reaction arrows to be negligible) we were able to further conclude that the KPL-IFF model is the simplest model able to explain all features.

The KPL-IFF model contains 5 parameters; kinetic proofreading (*k*_p_), limited signalling (*ϕ*), inhibition (*µ*), activation (*δ*), and amplification (*λ*). We used Approximate Bayesian Computations with Sequential Monte Carlo (ABC-SMC) to establish the set of these parameters that are able to reproduce the key features (44). We found that large variation of the parameters were possible provided that they obeyed certain relationships (see SI Appendix, Figure S6). For example, we found that *µ* and *δ* can vary by 1000-fold provided that *µ* > *δ* and that increases in *ϕ* can reproduce the phenotypic features provided that *k*_p_ decreased proportionally. Using the provided applet (see SI Appendix), we find that bell-shaped dose-response curves are less pronounced when the condition *µ* > δ is not satisfied. A large variation in the parameters is tolerated because the phenotypic features are scale-free (see SI Appendix).

### The KPL-IFF model predicts the T cell response to co-presentation of pMHC ligands

T cells generally experience mixtures of pMHC ligands when becoming activated. Previous studies have shown that co-presentation of an additional pMHC can modulate the T cell response in various ways, including both enhancing and inhibiting T cell activation (45).

To address the effects of pMHC co-presentation, we extended the KPL-IFF model to include an additional pMHC ligand with different binding kinetics and concentrations (Figure 5a). We used the extended KPL-IFF model to predict the T cell response to a titration of a lower affinity ligand in the presence of fixed concentrations of a higher affinity ligand (Figure 5b). As a result of the incoherent feedforward loop the model predicted a sigmoidal dose-response when the concentration of the high affinity ligand was left of its peak (≲.0.025) and a constant response when the concentration of the high affinity ligand was right of its peak (≳0.025). This is a direct result of the saturating activating pathway of the incoherent feedforward. Surprisingly, the model predicted that T cell activation cannot be inhibited by signals induced by the low affinity ligand even when the high affinity ligand is presented at concentrations that saturate the activation pathway of the incoherent feedforward. We confirmed these predictions by stimulating T cells with a titration of the lower affinity ligand, 5P, in the presence of fixed concentrations of the higher affinity ligand, 4A (Figure 5c). As predicted by the model, the dose-response curves appeared sigmoidal at lower doses of 4A and largely constant at higher doses, without any obvious inhibition of T cell activation by 5P.

**Figure 5:**
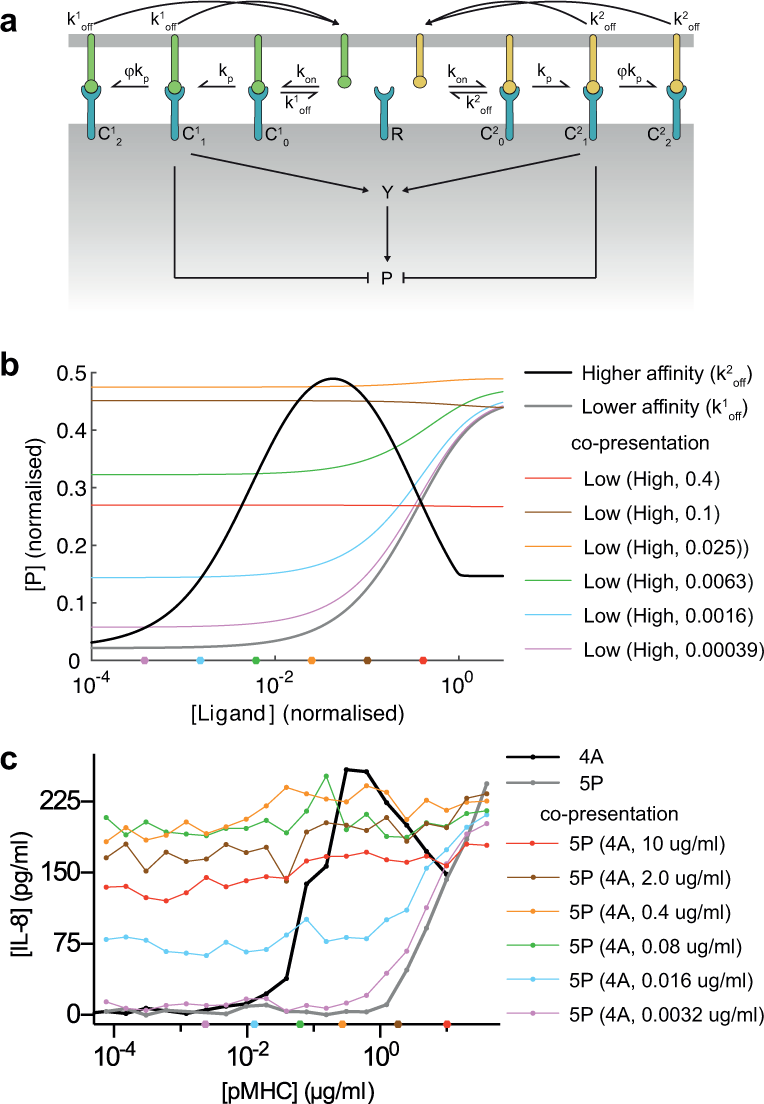
The KPL-IFF model predicts T cell activation in response to co-presentation of pMHC ligands. a) Schematic of signal integration by two distinct populations of pMHC ligands in the context of the KPL-IFF model. b) The model predicts that a titration of a low affinity ligand (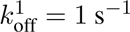) in the presence of a fixed concentration of a high affinity ligand (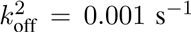) will be either sigmoidal or constant when the concentration of the high affinity ligand is left of its peak (purple, cyan, green) or right of its peak (orange, brown, red), respectively. Appreciable inhibition by the low affinty ligand is not predicted even when the activating pathway has saturated. c) T cell activation as measured by supernatant IL-8 released by Jurkat T cells in response to a titration of 5P (lower affinity ligand) at the indicated fixed concentrations of 4A (higher affinity ligand). The fixed concentration of the higher affinity ligand is indicated and labelled on the x-axis as coloured circles. Data are representative of 2 independent experiments. See SI Appendix for computational details.

## Discussion

We have measured the T cell response to a 1,000,000-fold variation in antigen affinity and dose. We found bell-shaped dose-response curves with a different pMHC (and hence different affinity) producing the largest T cell response at different doses. We show, without making prior biochemical assumptions and with the constraint of parsimony, that kinetic proofreading with limited signalling coupled to an incoherent feedforward loop (KPLIFF) is the only model that can explain all phenotypic features of the experimental data. We further confirmed predictions of the model concerning pMHC co-presentation. Remarkably, the KPL-IFF model can explain the T cell response to a 1,000,000-fold variation in antigen affinity and dose based on a simple pathway architecture despite the enormous molecular complexity in T cell signalling.

The present work has uncovered two independent mechanisms that lead to an optimal pMHC affinity. At low doses (left of the peak) we find that limited signalling through the TCR allows a single intermediate affinity pMHC to dominate the dose-response curve whereas at higher doses (right of the peak) a different pMHC affinity produces the most cytokine as a result of the bell-shaped dose-response curves produced by the incoherent feedforward loop. In light of our comprehensive data, it is likely that discrepancies between previous studies were a result of a limited range of tested pMHC affinity and dose. The model may account for previous work showing a bell-shape dose-response in the induction of regulatory T cells (5).

Modified TCRs and CARs often target tumour associated antigens that are differentially expressed between normal and cancer cells. Therefore, the antigen dose can be a critical determinant of succesful immunotherapy. As a result of the bell-shaped dose-response, we find that low affinity receptors can actually outperform high affinity receptors at high antigen doses. Our model provides a rationale for optimising the affinity of therapeutic receptors based on the target antigen dose, as recently proposed for a CAR (30). We provide a tool that can be used to examine the predicted T cell response for antigens of different affinities presented at different doses (Applet S1).

A number of studies have implicated negative feedback in TCR signalling (6), but we find that negative feedback cannot explain the phenotypic features of T cell activation. For example, negative feedback cannot produce bell-shaped dose-response curves. We note that our model does not preclude the existence of signalling proteins with negative effects, such as tyrosine phosphatases that can determine, for example, the net rate of TCR phosphorylation (*k*_*p*_). Negative feedback may be more important for the short timescale process of antigen discrimination rather than the longer timescale process of T cell activation that has been the focus of the present study (1, 6).

The limited signalling mechanism is related to previous work showing that a trade-off between serial binding and kinetic proofreading leads to an optimal pMHC half-life (10, 46). Serial binding of a single pMHC to many TCRs can increase signalling when the pMHC concentration is low and individual TCRs signal for a limited period of time upon binding. Under these conditions longer binding half-lives can reduce the number of productive TCR engagements (9). We find that at low doses reduced signalling is only observed when the TCR/pMHC half-life measured in solution is longer than 1 minute (e.g. low dose of 4A, 5Y, 8S compared to 9V in SI Appendix, Figure S2b,c). It follows that while limited signalling (and hence serial binding) may not be critical for physiological TCR-pMHC interactions, which have half-lives that last seconds, it is likely to be important for the design of high affinity therapeutic TCRs or CARs for T cell adoptive transfer therapies (25).

The internalisation of TCR is known to take place upon TCR triggering (1, 46) and it can be realised by different mechanisms. Intracellular signalling induced by activated TCR that leads to TCR internalisation is a form of negative feedback and, as discussed above, negative feedback cannot explain the observed phenotypic features. Limited signalling may result from the tagging of TCR for internalisation and explicitly including this internalisation, without incoherent feedforward, does not lead to bell-shaped dose-response curves in the steady-state (SI Appendix, Figure S8). The KPL-IFF model may implicitly be capturing TCR surface dynamics because directed movement (47) combined with polarised recycling (48, 49) of TCR into the immune synpase may balance with TCR internalisation (46) to maintain the relatively constant TCR concentration at the immune synapse assumed by the KPL-IFF model, which is consistent with previous calculations (50). A recent study has shown that changing the pMHC affinity can induce a programme that over a timescale of several days changes TCR levels (51). The KPL-IFF model can explain their observation that the higher affinity ligand induced greater TCR downregulation if TCR levels are determined by the output of the KPL-IFF model.

The systematic analyses revealed that a large number of more complex models can explain the phenotypic features (Figure 4e). This illustrates the broad challenge of 1) formulating unique models based on the known biochemistry and 2) relating the unique model we have formulated to the known biochemistry. Limited signalling may result from modification of the TCR signalosome, such as ubiquitination (52) and/or its movement into membrane environments incompatible with signalling (e.g. endosomes (46) or microvesicles (53)) (SI Appendix, Figure S7b). Kinetic proofreading can be realised by a number of different molecular mechanisms, such as sequential or random phosphorylation of the TCR (54, 55) and/or the recruitment of Lck associated coreceptors (56) (SI Appendix, Figure S7a). The incoherent feedforward loop may result from the fact that LAT can both activate (via Grb2 and SOS) and inhibit (via Dok1/Dok2 and RasGAP) Ras (7) (SI Appendix, Figure S7c) or from the observation that the TCR signalosome, by virtue of being able to associate with both a tyrosine kinase (ZAP-70) and a tyrosine phosphatase (SHP-1), can produce incoherent signals (2). Additionally, the positive and negative arms of the incoherent feedforward may represent two pathways that converge to regulate cytokine production. Future work is required to map the known biochemistry onto the KPL-IFF architecture.

The systematic search for parsimonious models that can reproduce phenotypic features of cellular activation, without prior biochemical assumptions, produces signalling pathways with tractable architectures. Just as subatomic details (e.g. nuclear structure) are not necessary for atomic molecular dynamics simulations, we argue that the correct description of signalling pathways may not require detailed biochemical knowledge of individual proteins. These predictive pathway models provide a mechanistic understanding of the modular network components required to integrate input signals from the cell surface into cellular activation outputs. Although they do not include full molecular detail, they offer an intuitive framework upon which biochemical information can be mapped, which has so far been elusive with reductionist approaches.

## Materials & Methods

### Protein production and surface plasmon resonance

HLA-A*02:01 heavy chain (residues 1-278) with C-terminal BirA tag and *β*_2_-microglobulin were expressed as inclusion bodies in E.coli, refolded in vitro in the presence of the relevant NY-ESO-1156−165 peptide variants (Table S1), and purified using size-exclusion chromatography. All peptides were purchased at >95% purity (Genscript, USA). Purified pMHC was biotinylated in vitro by BirA enzyme (Avidity, USA). The *α* and *β* subunits of the c58c61 (Clone 113) high affinity 1G4 T cell receptor (27) were expressed in E.coli as inclusion bodies, refolded in vitro, and purified using size exclusion chromatography as described previously (17).

TCR-pMHC binding affinity and kinetics were measured by surface plasmon resonance using a Biacore 3000 (GE Healthcare, USA) as previously described (17). Briefly, biotinylated pMHC were coupled to the CM5 surface by covalently coupled streptavidin with a target immobilisation level of 250 response units (RU) to minimise mass transport effects. The TCR analyte was diluted in HBS-EP running buffer and injected over the surface at 37°C using a flow rate of 30 µl/min. Running buffer was injected for 4 hours prior to the TCR injection when measuring interaction that relies on a long dissociation phase (i.e. high affinity interactions) to ensure that baseline drifts were minimal.

The off-rate (*k*_off_) was determined by fitting a one-phase exponential decay to the dissociation trace,

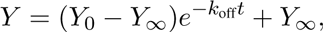

where *Y*_0_ and *Y*_∞_ are the initial and long-time asymptotic RU, respectively. The mean *k*_off_ across concentrations was used to determine *k*_on_. When the kinetics were such that the association phase could be resolved in time (i.e. sufficiently slow *k*_off_) we fit the following one-phase exponential association to the association trace,

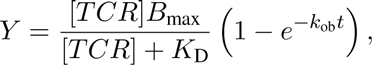

where *k*_ob_ = *k*_on_[TCR] + *k*_off_. We note that for high affinity interactions where the dissociation trace lifetime was > 15 minutes only a single concentration of TCR was used. Injection of multiple TCR concentrations is possible using the single-cycle kinetic mode but we found that these produced several incomplete association traces resulting in larger variability in *k*_on_ between experiments. We note that multiple analyte concentrations are particularly critical to determine the stoichiometry of the interaction. When the kinetics were such that the association phase could not be resolved in time (i.e. fast *k*_off_) we fit the following Langmuir binding equation to the steady-state response units in order to obtain an estimate for *K*_D_,

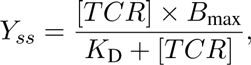

where *Y*_*ss*_ is the steady-state RU. The on-rate is determined using *k*_on_ = *k*_off_/*K*_*D*_. All data fitting was performed in Prism (Graphpad, USA).

### Production of lentivirus for transduction

HEK 293T cells were seeded into 175 cm^2^ flasks 24h prior to transfection to achieve 50-80% confluency on the day of transfection. Cells were co-transfected with the respective 3rd generation lentiviral transfer vectors and packaging plasmids using a standard PEI (polyethylenimine) transfection protocol as follows. The medium was replaced with serum-free DMEM. Transfer vector and the packaging plasmid mix (17.5 µg of pRSV-rev and pMDLg/pRRE as well as 6.8 *µ*g of pVSV-G) were diluted in 400 *µ*l of serum-free DMEM and a dilution of 112 *µ*g PEI in serum-free DMEM was prepared in another tube. Both were mixed vigorously and incubated at room temperature for 20 minutes. The mixture was added dropwise to the cells, which were then incubated at 37 °C/ 10% CO_2_ for 4-5 hours. Afterwards, the medium was replaced with complete medium. The supernatant was harvested and filtered through a 0.45 *µ*m cellulose acetate filter 24 hours later. Lentiviral particles were concentrated using Lentipac^TM^ Lentivirus concentrator (GeneCopoeia, USA) according to manufacturer’s protocol.

### Transduction of Jurkat T cells

The Jurkat E6.1 T cell line expressing the NFAT/AP-1 luciferase reporter and CD8*α* (57) were transduced with the c58c61 TCR. To do this, 3 million cells were resuspended in 2 mL of concentrated virus followed by centrifugation at 2095×g for 1-2 hours. The cells were incubated at 32 °C for 3.5-6 hours and then cultured at 37°C/ 10% CO_2_ in DMEM supplemented with 10% FBS, 100 U/ml Penicillin and 100 *µ*g/ml Streptomycin.

### Isolation and transduction of primary T cells

Peripheral blood mononuclear cells (PBMCs) were isolated from healthy donor blood by density gradient centrifugation: Blood collected in heparinised tubes was diluted 1:2 with PBS, carefully layered onto Ficoll-Paque^®^ in 50ml tubes and spun without brake at 400×g at room temperature for 30 minutes. The PBMCs were collected from the interphase, spun at 520×g for 5 minutes and washed once with PBS.

CD8+ T cells were isolated from PBMCs using the Dynabeads^®^ Untouched^TM^ Human CD8 T Cells kit (Life Technologies^TM^, USA) following manufacturer’s instructions. Briefly, PBMCs were resuspended in Isolation Buffer (0.1% BSA and 2 mM EDTA in PBS), blocked with FBS and unwanted cells were labelled with an antibody mix (containing biotinylated antibodies for human CD4, CD14, CD16, CD19, CD36, CD56, CDw123 and CD235a). Subsequently, the PBMCs were washed and incubated with streptavidin-coated Dynabeads ®. The suspension was resuspended thoroughly with Isolation Buffer before the tube was placed into a magnet. The supernatant containing ‘untouched’ CD8^+^ T cells was collected. This process was repeated twice and the supernatants were combined.

The isolated CD8+ T cells were spun at 520×g for 5 minutes and resuspended at a concentration of 10^6^ cells/ml in completely reconstituted DMEM, supplemented with 50 units/ml IL-2 and 10^6^ CD3/CD28-coated Human TActivator Dynabeads^®^ (Life Technologies) per ml. Cells were cultured at 37°C/ 10% CO_2_ overnight.

The next day, 10^6^ purified primary human CD8^+^ T cells in 1 ml of medium were transduced with 1 ml of concentrated virus supplemented with 50 units of IL-2. The cells were cultured at 37°C/ 10% CO_2_ and the medium was replaced with fresh medium containing 50 units/ml IL-2 every 2-3 days. CD3/CD28-coated Dynabeads^®^ were removed on day 5 after lentiviral transduction and the cells were characterised and used for experiments once the populations expanded to adequate sizes.

### T cell stimulation

Streptavidin-coated 96-well plates (Sigma-Aldrich, USA) were washed 2× with PBS 0.05% tween followed by 1× with PBS. Plates were incubated at 37°C with PBS 1% BSA for 1 hour. Serially diluted pMHC (in PBS) were transferred to the plates and incubated at 4°C for 90 minutes (volume of 100 *µ*l per well). Plates were washed 3× with PBS following incubation. Plates were always prepared in pairs so that one plate could be used for the stimulation assay and the other to determine the levels of correctly folded plate-immobilized pMHC.

T cell stimulation assays were performed by first washing and resuspending the cells in culture media without IL-2. T cells were then added at 50,000 cells per well in a volume of 100 *µ*l. Plates were spun at 9×g (4 min) and then incubated at 37°C/ 10% CO_2_ for the required stimulation time.

Concentrations of supernatant cytokines were determined using commercially available ELISA kits following manufacturers’ protocols: OptEIA IFN-*γ* (BD Biosciences, USA, 555142) and 2nd Generation Ready-Set Go! Kits (Ebioscience, USA) for MIP-1*β* (88-7034-88) and IL-8 (88-8086-88). Measurement of AP1/NFAT activity in Jurkats was performed by lysing cells using ONE-GloTM Luciferase substrate (Promega, USA, E6110) for 5 minutes before luminescence was read using a PherastarPlus plate reader (BMG Lab Tec, Germany). Data were corrected for background luminescence using unstimulated cells.

Levels of active plate-immobilized pMHC were measured on the second plate using mouse anti-human HLA class I antibody (Clone W6/32; Ebioscience, 14-9983) in combination with fluorescent secondary goat anti-mouse IgG IRDye 800CW antibody (LI-COR, USA, 926-32210). Fluorescence measurements were performed with the Odyssey Imaging system (LI-COR). A Hill function was fit to the fluorescence over the initial pMHC concentration (in *µ*g/ml) to determine the *EC*_50_ for each pMHC using Prism. The pMHC concentrations in the functional assays were modified to reflect differences in the immobilisation *EC*_50_ as follows: log[pMHC]^corrected^ = log[pMHC] + (log(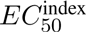) - log(*EC*_50_)) where the 9V pMHC ligand served as the index.

### Flow cytometery assays

#### Intracellular cytokine staining

T cell stimulation was performed as described above except that 50 *µ*l of medium supplemented with Brefeldin A (2.5 *µ*g/ml final concentration) was added to the respective samples followed by spinning at 520×g for 5 minutes before returning the cells to the incubator for another 2 hours (primary T cells, 4 hours total stimulation) or for another 3 hours (Jurkat T cells, 8 hours total stimulation).

After stimulation, cells were spun at 520×g for 5 minutes and resuspended with 2 mM EDTA in PBS. After an additional spin, the cell pellets were fixed by resuspension in 50 *µ*l/well 4% formaldehyde in cold PBS at 4°C (10 min). Fixed cells were washed with PBS, spun down at 520×g for 10 minutes and resuspended in 100 *µ*l/well permeabilisation buffer (PBS with 2% BSA and 0.1% TritonX100). After 10 minutes at 4°C, cells were spun at 520×g for 10 minutes, and resuspended in permeabilisation buffer containing the respective antibody (E8N1 APC-conjugated IL-8 antibody or 4S.B3 AlexaFluro647 conjugated IFN-_ antibody, BioLegend) for 20 minutes at 4°C. After washing twice with PBS (520×g for 10 min), cells were resuspended in 100-150 **µ**l PBS per sample and transferred into FACS tubes for analysis. Jurkat cells had to be spun before the transfer to mitigate cell losses.

#### Annexin V assay

Jurkat T cells were stimulated as described above except that 100,000 cells were used per well. After 16 hours of stimulation, cells were removed first by gently pipetting them out of each well and second by washing each well with PBS. Cells were transferred to 1.5 ml tubes and washed 2× with PBS. Cells were resuspended in Annexin-V buffer (HEPES pH 7.4 10 mM, NaCl 140 mM, CaCl2 2.5 mM) at a concentration of 1-5×10^6^ cells/ml. Cells were stained with PE-Annexin-V (BD Biosciences, 556421) at a concentration of 5 **µ**l per 100 **µ**l cells and incubated at room temperature in the dark for 15 minutes. After washing twice with PBS, samples were ready for flow cytometry.

#### c58c61 T cell receptor expression

5x10^5^ T cells per sample were washed with PBS in FACS tubes (3 ml, 5 min at 520×g) and stained with high-affinity 9V pMHC (7 **µ**g/ml; 200**µ**l/sample) for 30min. Subsequently, they were washed with PBS and stained with R-PE-conjugated streptavidin (AbD Serotec, USA, STAR4A, 1:100; 200 **µ**l/sample) for another 30 min. After washing twice with PBS, samples were ready for flow cytometry.

All flow cytometry was performed using a FACSCalibur (BD Biosciences) with at least 10,000 cells. All analysis was performed using the software FlowJo (Treestar, USA).

## Author contributions

ML, HSL, PK, JN, NT, EAS performed experiments; ML, PK, NT, and OD performed the mathematical modelling; ML, HSL, PK, JN, NT, EAS, PKM, PAvdM, OD analysed data; ML, HSL, PK, PKM, PAvdM, OD designed the research and wrote the paper; ML, HSL, and PK contributed equally to the study. All authors discussed the results and commented on the paper.

## Acknowledgements

We thank Gillian Griffiths, Paul Francois, Arup Chakraborty, John R. James, and Bela Novak for feedback during the development of the work. We acknowledge Georg Kempf, Marcus Bridge, Nathan Jenko, Ann Tivey, and Dhruv Jayanth for assistance in protein production and kinetic measurements. We thank Simon J. Davis for a critical reading of the manuscript. The c58c61 TCR was provided by Adaptimmune Ltd. ML and NT are supported by a Doctoral Training Centre Systems Biology studentship from the Engineering and Physical Sciences Research Council (EPSRC). PAvdM is funded by a Wellcome Trust Senior Investigator Award (Grant Number: 101799). The work is funded by a Sir Henry Dale Fellowship (to OD) jointly funded by the Wellcome Trust and the Royal Society (Grant Number: 098363).

## References

1. Smith-Garvin JE, Koretzky Ga, Jordan MS (2009) T cell activation. Annual review of immunology 27: 591–619.

2 Altan-Bonnet G, Germain RN (2005) Modeling T cell antigen discrimination based on feedback control of digital ERK responses. PLoS biology 3:e356.

3. Dushek O, van der Merwe PA (2014) An induced rebinding model of antigen discrimination. Trends in Immunology 35: 153–158.

4. Turner MS, Kane LP, Morel Pa (2009) Dominant role of antigen dose in CD4+ Foxp3+ regulatory T cell induction and expansion. The Journal of Immunology 183: 4895–4903.

5. Gottschalk Ra, Corse E, Allison JP (2010) TCR ligand density and affinity determine peripheral induction of Foxp3 in vivo. The Journal of experimental medicine 207: 1701–1711.

6. Acuto O, Di Bartolo V, Michel F (2008) Tailoring T-cell receptor signals by proximal negative feedback mechanisms. Nature Reviews Immunology 8: 699–712.

7. Brownlie RJ, Zamoyska R (2013) T cell receptor signalling networks: branched, diversified and bounded. Nature Reviews Immunology 13: 257–269.

8. Corse E, Gottschalk Ra, Allison JP (2011) Strength of TCR-peptide/MHC interactions and in vivo T cell responses. Journal of immunology 186: 5039–5045.

9. Lever M, Maini PK, van der Merwe PA, Dushek O (2014) Phenotypic models of T cell activation. Nature Reviews Immunology 14: 619–629.

10. Kalergis aM, et al. (2001) Efficient T cell activation requires an optimal dwell-time of interaction between the TCR and the pMHC complex. Nature immunology 2: 229–234.

11. Coombs D, Kalergis AM, Nathenson SG, Wofsy C, Goldstein B (2002) Activated TCRs remain marked for internalization after dissociation from pMHC. Nature immunology 3: 926–931.

12. Holler PD, Kranz DM (2003) Quantitative analysis of the contribution of TCR/pepMHC affinity and CD8 to T cell activation. Immunity 18: 255–264.

13. Gonz´alez Pa, et al. (2005) T cell receptor binding kinetics required for T cell activation depend on the density of cognate ligand on the antigen-presenting cell. Proceedings of the National Academy of Sciences of the United States of America 102: 4824–4829.

14. McMahan RH, et al. (2006) Relating TCR-peptide-MHC affinity to immunogenicity for the design of tumor vaccines. Journal of Clinical Investigation 116: 2543–2551.

15. Chervin AS, et al. (2009) The impact of TCR-binding properties and antigen presentation format on T cell responsiveness. Journal of immunology 183: 1166–1178.

16. Schmid Da, et al. (2010) Evidence for a TCR affinity threshold delimiting maximal CD8 T cell function. Journal of immunology (Baltimore, Md.: 1950) 184: 4936–4946.

17. Aleksic M, et al. (2010) Dependence of T cell antigen recognition on T cell receptor-peptide MHC confinement time. Immunity 32: 163–174.

18. Govern CC, Paczosa MK, Chakraborty AK, Huseby ES (2010) Fast on-rates allow short dwell time ligands to activate T cells. Proceedings of the National Academy of Sciences 107: 8724–8729.

19. Corse E, Gottschalk Ra, Krogsgaard M, Allison JP (2010) Attenuated T cell responses to a high-potency ligand in vivo. PLOS BIOLOGY 8: 1–12.

20. Thomas S, et al. (2011) Human T cells expressing affinity-matured TCR display accelerated responses but fail to recognize low density of MHC-peptide antigen. Blood 118: 319–329.

21 Dushek O, et al. (2011) Antigen potency and maximal efficacy reveal a mechanism of efficient T cell activation. Science Signaling 4:ra39.

22. Irving M, et al. (2012) Interplay between T cell receptor binding kinetics and the level of cognate peptide presented by major histocompatibility complexes governs CD8+ T cell responsiveness. The Journal of biological chemistry 287: 23068–23078.

23. Lee HM, Bautista JL, Scott-Browne J, Mohan JF, Hsieh CS (2012) A Broad Range of Self-Reactivity Drives Thymic Regulatory T Cell Selection to Limit Responses to Self. Immunity 37: 1–12.

24. Tan MP, et al. (2015) T cell receptor binding affinity governs the functional profile of cancer-specific CD8+ T cells. Clinical & Experimental Immunology 180: 255–270.

25. Restifo NP, Dudley ME, Rosenberg Sa (2012) Adoptive immunotherapy for cancer: harnessing the T cell response. Nature Reviews Immunology 12: 269–281.

26. Jakka G, et al. (2013) Antigen-specific in vitro expansion of functional redirected NY-ESO-1-specific human CD8+ T-cells in a cell-free system. Anticancer Research 33: 4189–4202.

27. Li Y, et al. (2005) Directed evolution of human T-cell receptors with picomolar affinities by phage display. Nature Biotechnology 23: 349–354.

28. Chmielewski M, Hombach A, Heuser C, Adams GP, Abken H (2004) T Cell Activation by Antibody-Like Immunoreceptors: Increase in Affinity of the Single-Chain Fragment Domain above Threshold Does Not Increase T Cell Activation against Antigen-Positive Target Cells but Decreases Selectivity. The Journal of Immunology 173: 7647–7653.

29. Haso W, et al. (2012) Anti-CD22-chimeric antigen receptors targeting B cell precursor acute lymphoblastic leukemia. Blood 121: 1165–1174.

30. Caruso HG, et al. (2015) Tuning sensitivity of CAR to EGFR density limits recognition of normal tissue while maintaining potent antitumor activity. Cancer Research 75: 3505–3518.

31 Sadelain M, Brentjens R, Rivi`ere I (2009) The promise and potential pitfalls of chimeric antigen receptors. Current opinion in immunology 21.

32. Aldridge BB, Burke JM, Lauffenburger Da, Sorger PK (2006) Physicochemical modelling of cell signalling pathways. Nature cell biology 8: 1195–1203.

33 Gunawardena J (2014) Models in biology: ‘accurate descriptions of our pathetic thinking’. BMC Biology 12:29.

34. Schmidt M, Lipson H (2009) Distilling free-form natural laws from experimental data. Science 324: 81–85.

35. Ma W, Trusina A, El-Samad H, Lim Wa, Tang C (2009) Defining network topologies that can achieve biochemical adaptation. Cell 138: 760–773.

36 Shah Na, Sarkar Ca (2011) Robust network topologies for generating switch-like cellular responses. PLoS Computational Biology 7.

37. Francçois P (2014) Evolving phenotypic networks in silico. Seminars in cell & developmental biology 35: 6–13.

38 Villaverde AF, Banga JR (2014) Reverse engineering and identification in systems biology: Strategies, perspectives and challenges. Journal of the Royal Society Interface 11.

39 Daniels BC, Nemenman I (2014) Automated adaptive inference of coarse-grained dynamical models in systems biology. Nature Communications 6:38.

40. Sami M, et al. (2007) Crystal structures of high affinity human T-cell receptors bound to peptide major histocompatibility complex reveal native diagonal binding geometry. Protein engineering, design & selection: PEDS 20: 397–403.

41 Adams JJ, et al. (2015) Structural interplay between germline interactions and adaptive recognition determines the bandwidth of TCR-peptide-MHC cross-reactivity. Nature Immunology 17.

42. Green DR, Droin N, Pinkoski M (2003) Activation-induced cell death in T cells. Immunological reviews 193: 70–81.

43. Alon U (2007) Network motifs: theory and experimental approaches. Nature Reviews Genetics 8: 450–461.

44. Toni T, Welch D, Strelkowa N, Ipsen a, Stumpf MP (2009) Approximate Bayesian computation scheme for parameter inference and model selection in dynamical systems. Journal of The Royal Society Interface 6: 187–202.

45. Krogsgaard M, Juang J, Davis MM (2007) A role for “self” in T-cell activation. Seminars in immunology 19: 236–244.

46 Valitutti S, Muller S, Cella M, Padovan E, Lanzavecchia A (1995) Serial triggering of many T-cell receptors by a few peptide MHC complexes. Nature 375.

47. Moss WC, Irvine DJ, Davis MM, Krummel MF (2002) Quantifying signaling-induced reorientation of T cell receptors during immunological synapse formation. Proc Natl Acad Sci U S A 99: 15024–15029.

48. Das V, et al. (2004) Activation-induced polarized recycling targets T cell antigen receptors to the immunological synapse; involvement of SNARE complexes. Immunity 20: 577–588.

49. Finetti F, et al. (2014) Specific recycling receptors are targeted to the immune synapse by the intraflagellar transport system. Journal of cell science 127: 1924–1937.

50. Arkhipov SN, Maly IV (2006) Quantitative analysis of the role of receptor recycling in T cell polarization. Biophysical journal 91: 4306–4316.

51 Gallegos AM, et al. (2016) Control of T cell antigen reactivity via programmed TCR downregulation. Nature immunology 1.

52. Yang M, et al. (2015) K33-linked polyubiquitination of Zap70 by Nrdp1 controls CD8+ T cell activation. Nature Immunology 16: 1–12.

53. Choudhuri K, et al. (2014) Polarized release of T-cell-receptor-enriched microvesicles at the immunological synapse. Nature 507: 118–123.

54. McKeithan TW (1995) Kinetic proofreading in T-cell receptor signal transduction. Proceedings of the National Academy of Sciences of the United States of America 92: 5042–5046.

55. Mukhopadhyay H, et al. (2016) Multisite phosphorylation of the T cell receptor z-chain modulates potency but not the switch-like response. Biophys. J. 110: 1896–1906.

56. Stepanek O, et al. (2014) Coreceptor Scanning by the T Cell Receptor Provides a Mechanism for T Cell Tolerance. Cell 159: 333–345.

57. Paster W, et al. (2015) A THEMIS:SHP1 complex promotes T-cell survival. The EMBO journal 34: 393–409.

